# Extracellular vesicle-mediated Transfer of miR-1 from Primary tumor Repress Distant Metastasis Growth

**DOI:** 10.1101/2023.05.19.541440

**Authors:** Hui-Su Kim, Kang-Hoon Lee, Keun Hong Son, Tae-Jin Shin, Je-Yoel Cho

## Abstract

Metastases originate from primary tumors that conquer distant organs. Growing evidence suggests that metastases are still under primary tumor dominance, even out of the primary territory. However, the mechanism by which primary tumors prime the fate of metastasis is unclear. Here, we investigated the influence of primary tumor-derived extracellular vesicles (pTDEs) on distant metastasis. Metastatic growth was remarkably suppressed when cells were treated with pTDEs *in vitro* and intravenously in a spontaneous metastatic mouse model. pTDEs from primary tumors with cancer stem cell (CSC) characteristics inhibit metastatic growth by inducing intracellular ROS/DNA double-stranded damage, thus inhibiting G2/M phase accumulation. We found that miR-1 was the most abundant in pTDEs and showed a stronger inhibitory effect when miR-1 was overloaded in pTDEs. Collectively, we demonstrated that the primary tumor could control distant metastatic cancer via primary tumor-derived EV-miR-1. Additionally, our approach suggests the development of anticancer drugs for metastasis based on pTDE-miR-1.

## Introduction

Primary tumors evict metastases to leave home, but keep them under control^1–3^. Many studies have revealed that primary tumors act on the tumor microenvironment to inhibit the growth of metastases, thereby interfering with angiogenesis or being eliminated by immune cells ^4–6^. Therefore, it is not surprising that in patients whose primary tumor has been excised, as spontaneous inhibition disappears, metastases that escape inhibition begin to grow and cause recurrence. Ancient Roman physician Celsus first observed that the carcinoma had returned after excision.^7^. A century ago, Tyzzer reported that the excision of the primary tumor resulted in larger metastases ^8^. From 1900 to the present, many studies have reported that the primary tumor restricted distant metastatic growth, and the growth of sudden metastases was observed when the primary tumor was resected ^9–13^. One fundamental finding is that angiostatin and endostatin interrupt angiogenesis of metastases and lead to starvation ^14, 15^. Furthermore, primary tumors can restrict metastases indirectly by remodeling CD8+ cytotoxic T cells and macrophages to eradicate them ^1, 3, 16^.

Since the discovery of extracellular vesicles, cells have been known to deliver their messages by wrapping them in a lipid bilayer to communicate with distant cells^17^. Extracellular vesicles (EVs) contain proteins, lipids, and nucleic acids and referred to as exosomes or microvesicles, depending on their size and biogenesis. Research on the biological function of EVs in cancer has mainly focused on their role in tumorigenesis and the formation of the tumor microenvironment ^18^, and there have been a few attempts to address the function of EVs as controllers of metastases ^19–22^. Despite numerous studies of tumor-derived EVs (TDEs), there is no evidence that pTDEs can metastasize.

In this study, we demonstrated that primary tumors used EVs to control metastatic growth. Moreover, EV miR-1 acts as a key regulator of inhibition, and engineered pTDEs for enhancing miR-1 intensify the inhibition function in both cell and mouse metastases. These findings highlight that the primary tumor sends messages through pTDEs in the regulation of metastases.

## Results

### EVs from primary tumors inhibit the growth of metastases

To investigate the role of primary tumor controls in metastases, we used cell lines of both primary and metastatic tumors established from the same spontaneous cancer patient. As there is no pair of naturally occurring human breast cancer (HBC)-derived primary and metastatic tumor cell lines, we used canine mammary gland tumors (CMT), which have recently been reported to have pathological and molecular characteristics similar to those of HBC ^23, 24^. The pair of CMT cell lines (primary/metastases, CHMp/CHMm) originated from the same patient with CMT.

We investigated whether primary tumor-derived EVs could inhibit the formation and growth of metastases. CHMp- and CHMm-derived EVs (pTDEs and mTDEs) were isolated as described in the Methods and Extended Data Fig. 1a. Purified EVs had an average of 147 nm (pTDEs) and 143 nm (mTDEs) cup-shaped vesicles and were also positive for TSG101 and Alix (Extended Data Fig. 1b-d).

To establish a metastatic mouse model, we followed the surgical methods published by Piranlioglu et al.^3^ In brief, primary tumor was surgically removed 21 days after CHMp cell injection (Extended Data Fig. 2a-c). All 20 mice underwent surgery, including four (20%) sham surgeries. In mice that underwent sham surgery, the primary tumor continued to grow and finally exceeded 10 mm^3^ within two weeks, and unexpected intestinal invasion was observed instead of metastatic lung cancer (Extended Data Fig. 2d). Prior to injecting pTDEs, PKH67-labeled EVs were injected to determine their exact accumulation sites after intravenous administration. PKH67-labeled pTDEs were highly enriched in disseminated and primary tumors. (Extended Data Fig. 2e).

Subsequently, the mice were divided into two groups: one group was regularly injected with pTDEs and the other group with PBS as a control. The injection schedule is shown in Figure 1a. The bioluminescence image showed very little metastatic growth in the pTDE group compared to the control group on the 39^th^ day (Fig. 1b,c). Metastases occurred in three out of four mice in the control group, but only very minor signals were observed in two out of five mice injected with pTDEs (Fig. 1b-c). The lungs from control mice exhibited a higher number of metastatic nodules than those from pTDE-injected mice (Fig. 1d,e). To date, a mouse model has shown that removing primary tumors could promote the growth of metastatic cancers^25–28^, as seen in clinical findings. Thus, it is reasonable to infer that disseminated metastases were dormant in the lung metastatic site, waiting for a wake-up and growth signal, such as the absence of pTDEs. Here, we demonstrated that pTDEs play an important role in the inhibition of metastases.

**Figure 1.**
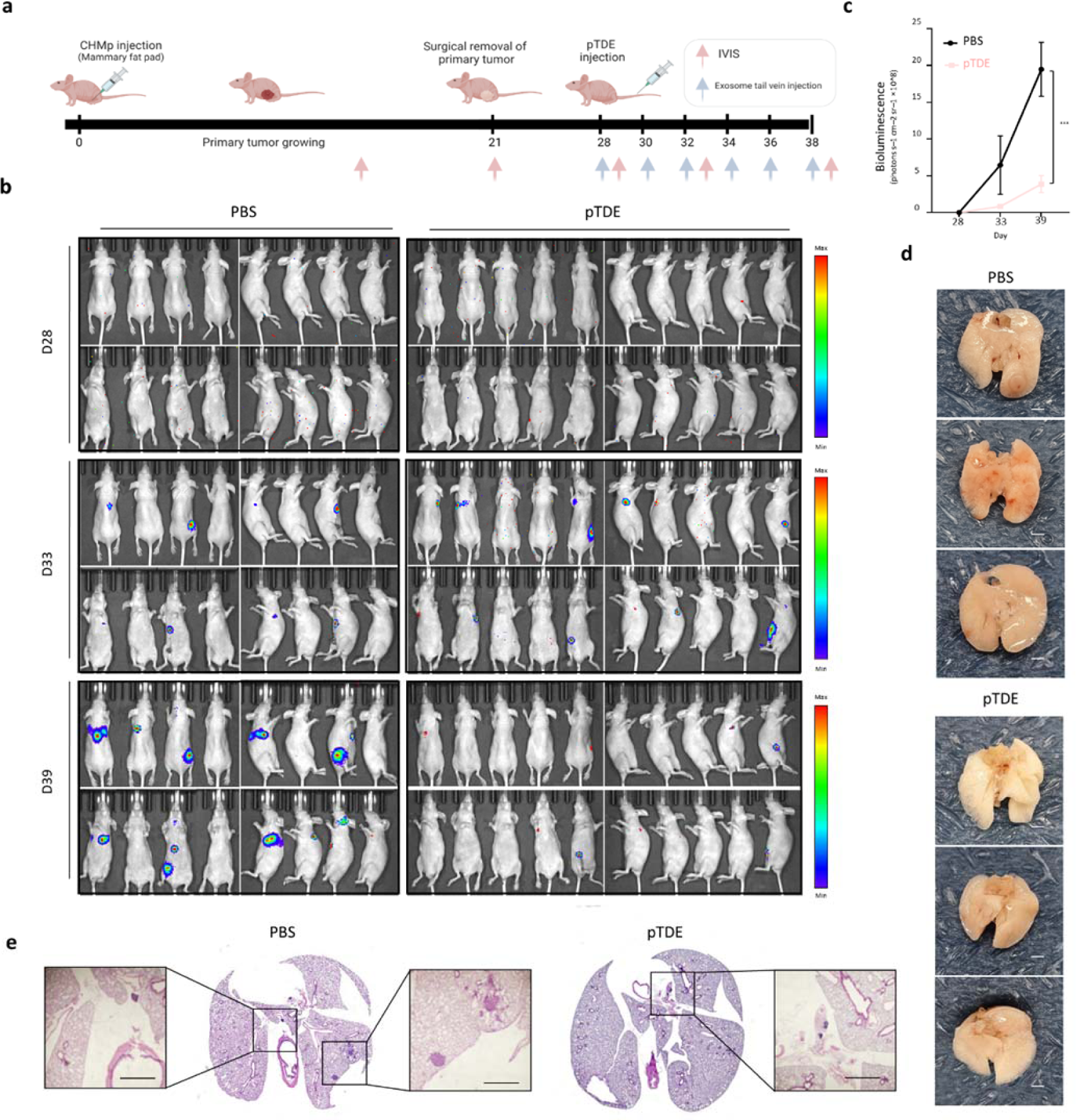
Primary tumor-derived EVs (pTDEs) restrict metastasis formation and growth. **(a)** Schematic illustration of the established experimental metastasis model. Mice were inoculated with CHMp cells into the mammary fat pad to produce a primary tumor on day 0 (n = 20). On day 21, primary tumors were surgically removed. Seven days after surgery, PBS (n = 4) and CHMp EVs (n = 5) were administered to mice. Mice were treated with PBS or pTDEs (10ug/mouse) through the tail vein with six times every two days. **(b)** Bioluminescence images of CHMp lung metastases after surgery for confirmation of residual primary tumor mass (at D28). D33 bioluminescence images were acquired from the mice treated with PBS or pTDEs three times, and D39 images were acquired six times at two-day intervals. For each group of daily images, four mouse images were acquired at four positions (dorsal, ventral, right lateral and left lateral) to capture every possible signal from the mice. **(c)** Graph showing quantification of lung metastases treated with PBS and pTDEs in a metastasis mouse model. Unpaired Student’s t test was used to compare groups. (\*\*\**p <* 0.001) **(d)** Picture of lungs after PBS and pTDE treatment. Top; PBS treated, bottom; pTDE treated. Scale bar, 2 mm. **(e)** Representative H&E staining of lung tissue (4X) in the metastasis model. Left for PBS-treated mice, right for pTDE-treated mice. Magnified images : Scale bar, 1 mm. Results are presented as mean ± SD (4-5 mice were used for each group)

### pTDEs autonomously suppress metastasis growth in a direct and indirect manner

To determine how pTDEs block metastasis, we treated CHMm metastatic cancer cells and endothelial cells with pTDEs. We used mTDE as another control for pTDE to rule out any influence of changes in the process of EV purification. CHMm cells treated with pTDEs exhibited reduced viability and proliferation, whereas mTDE was not significantly different from the PBS control (Fig. 2a, b). Moreover, pTDEs showed higher accumulation in the G2/M phase of the cell cycle than in the control cells (Fig. 2c). Since it is known that accumulation in the G2/M phase can be caused by cellular stress, such as cell oxidation, DNA replication, and transcription ^29, 30^ we investigated whether pTDE treatment elicited stress conditions. Treatment with pTDEs caused a marked increase in intracellular ROS levels (Fig. 2d) and the level of γ-H2A.X, a sensitive marker of DNA damage (Fig. 2e, f). These results indicate that pTDEs induce an increase in intracellular ROS, genomic instability, accumulation in the G2/M cell phase, and suppression of the growth of metastatic cells. However, this growth inhibition did not correlate with cell death or mobility (Extended Data Fig. 3a-c).

**Figure 2.**
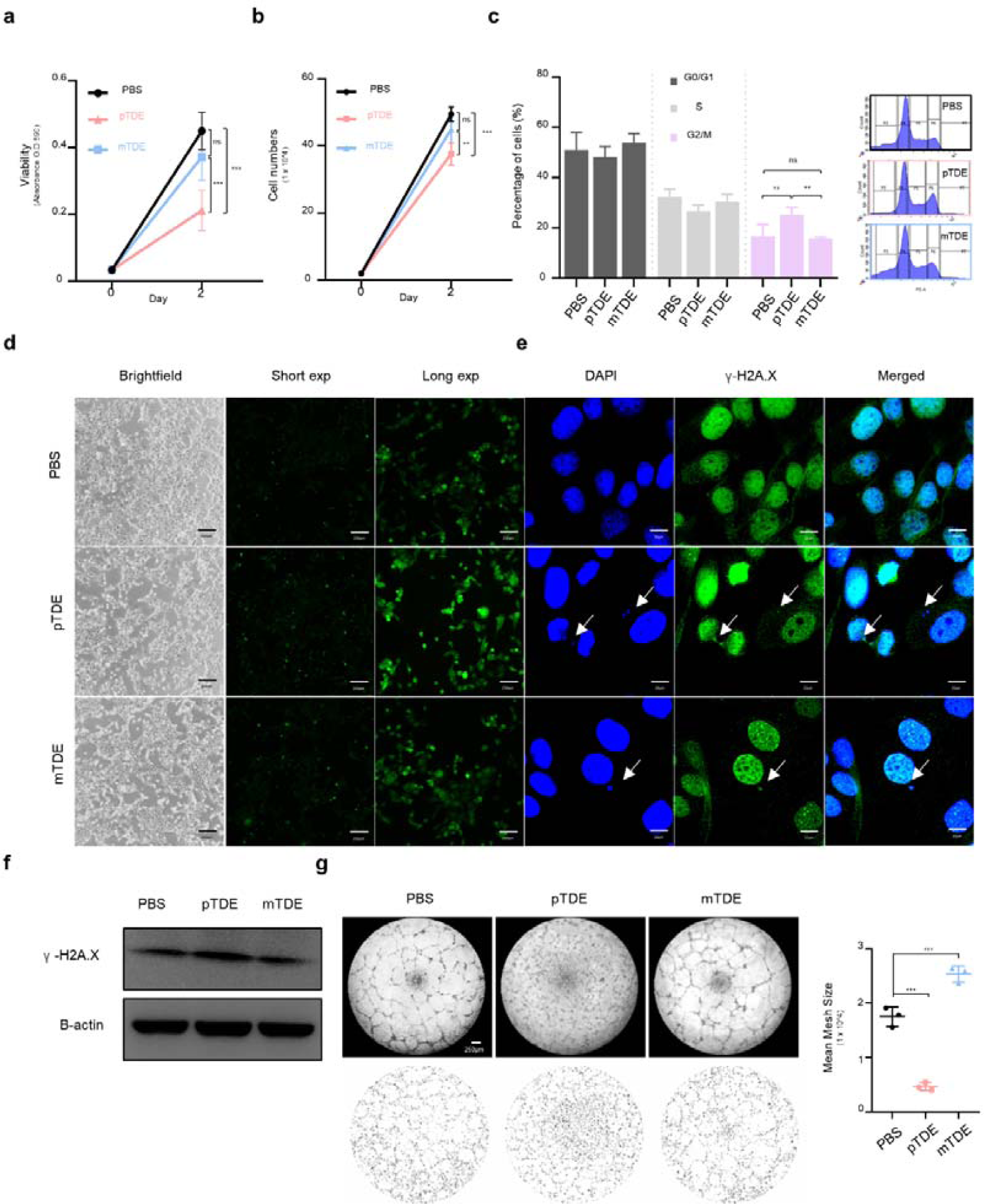
pTDEs induce cellular ROS, DNA damage, and G2/M arrest, resulting in inhibition of proliferation in recipient cells. **(a)** Recipient cell viability after tumor-derived EVe (TDE) treatment (0.1ug/ml) was measured by MTT assay (n=8). **(b)** The proliferation rate was manually measured by cell counting in the PBS and TDE treatment groups. pTDEs decreased cell viability and proliferation. **(c)** Cell cycle analysis of EV-treated cells was performed using flow cytometry. pTDE induced an increase in G2/M phase. **(d)** Representative images showing cellular ROS levels. Intracellular oxidative stress was measured with the fluorescent ROS probe H2DCFDA (2,7- dichlorodihydrofluorescein diacetate). Brighter green fluorescence indicates high ROS. EV-treated cells were stained with H2DCFDA to measure intracellular ROS. Scale bar, 170 μm **(e)** For measuring damaged DNA, γ-H2A.X (green) and DAPI (blue) were stained in EV-treated CHMm cells. The white arrow indicates damaged DNA, which colocalized with γ-H2A.X and DAPI. Scale bar, 10 μm. **(f)** Western blot for intracellular γ-H2A.X, γ-H2A.X was increased with pTDE treatment. **(g)** Tube formation assays of HUVECs to detect the angiogenesis potential of pTDEs. HUVECs were seeded on Matrigel, and the EVs were treated for 24 h. Representative images showing that pTDEs inhibited tube formation compared with controls. Bottom microscopic images were analyzed by the ‘ImageJ plugin Angiogenesis analyzer.’ Quantification of the mean mesh size is presented as the mean ± SEM. Two-way ANOVA and Tukey’ HSD test was used to compare groups; **P < 0.01, *** p < 0.001, and ns, not significant. Results are presented as mean ± SD.

In contrast, pTDEs had a strong effect on tube formation by endothelial cells (Fig. 2g). Human umbilical vein endothelial cells (HUVECs) incubated with pTDEs showed a 1.5-fold decrease in mesh area compared to that of PBS-treated cells, which formed regular tubes. These findings revealed that pTDE-treated mice showed restricted metastases by directly inhibiting metastatic tumor cell growth and indirectly manipulating endothelial cells of the tumor microenvironment by inhibiting angiogenesis.

### Primary tumor stem cell-derived EVs restrict metastases growth

To further investigate how primary tumors awaken their force to control metastases, we compared the stemness between primary and metastatic tumors (Extended Data Fig.4a). CD44+/CD24-expression, a representative marker of breast cancer stem cells (CSCs), showed a higher CSC ratio in the primary tumor than in metastases (Fig 3a). In addition, other breast CSC characteristics, such as fast cell proliferation, high aldehyde dehydrogenase (ALDH) enzyme activity, and mammosphere formation, were substantially more enriched in CHMp cells than in CHMm cells (Fig. 3b-e). Spheroids were fluorescently labeled with CD44 and CD24, and only the CSC populations formed spheroids (Fig. 3e). We also compared CD44, CD24, and ALDH1A1 protein levels and CSC-related gene expression in CHMp and CHMm cells (Extended Data Fig. 4b, c and Supplementary Table 1), including CD44, ALDH, drug resistance-related ABC transporters, stemness factors, and epithelial-mesenchymal markers. CHMp cells had higher levels of CD44, ALDH subtypes, ABCG2, and interestingly, mesenchymal characteristics than CHMm cells. Next, the CSC-rich CHMp cells formed a larger and faster tumor than CHMm cells and showed vast differences even when the same number of cells was inoculated (Fig. 3f-h and Extended Data Fig. 4d). These data suggest that primary tumor CHMp cells have more CSCs than CHMm cells do.

**Figure 3.**
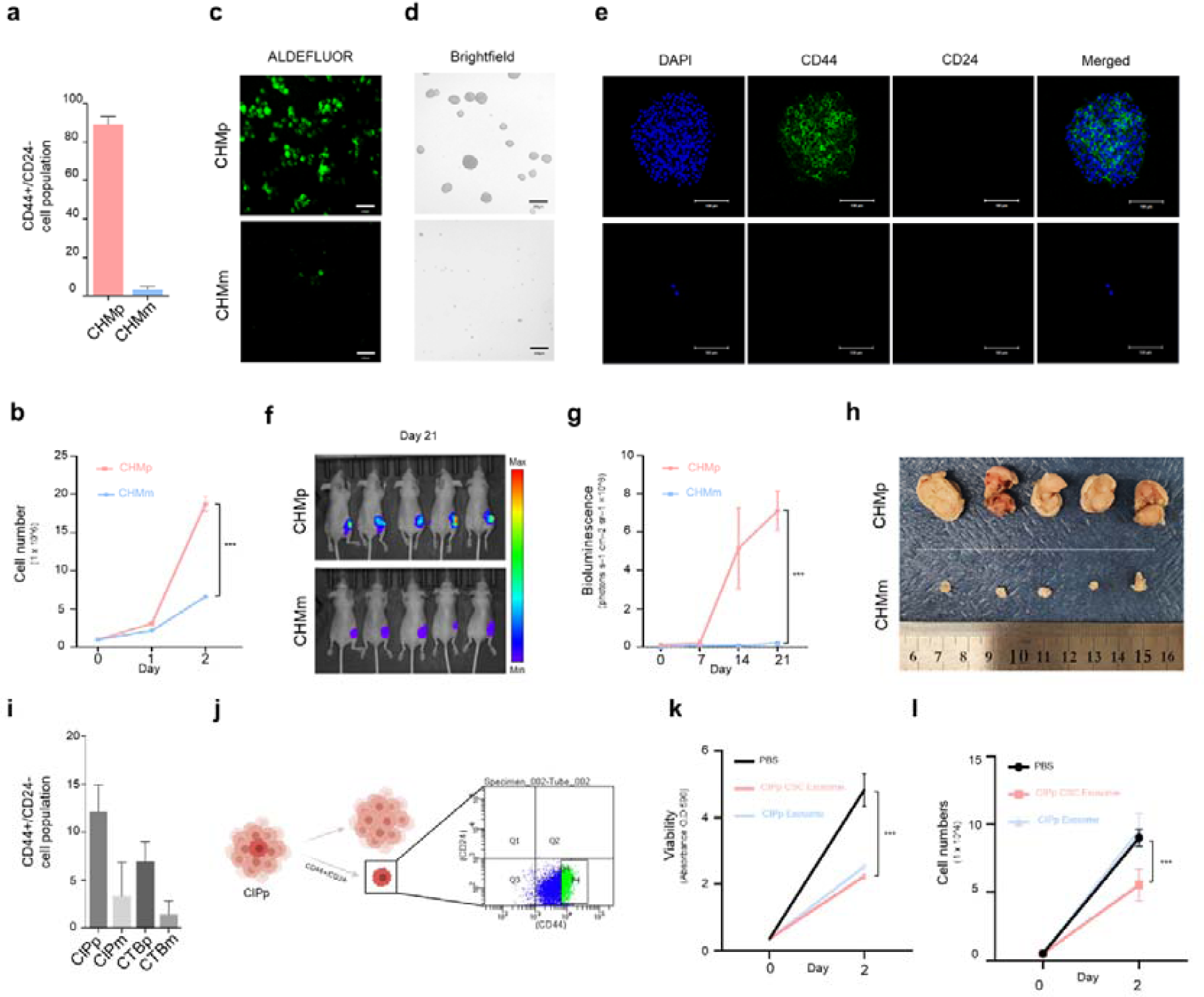
Cancer stemness contributes to the suppression of metastases. **(a)** CMT cell lines were stained with CD44 and CD24 antibodies and analyzed by flow cytometry. **(b)** The growth rate of CHMp and CHMm cells. Cell proliferation was measured by manual cell counting. **(c)** ALDH activity was examined using an ALDEFLUOR assay in CHMp and CHMm cells. Fluorescent images were examined using a 40x objective lens; the scale bar indicates 130 μm. **(d)** Representative confocal microscopy images of mammospheres formed by CHMp and CHMm. Scale bar, 400 μm. **(e)** Mammospheres were stained with antibodies against the CSC markers CD44 (green) and CD24 (red). DAPI (blue) was used as a nuclear marker. Scale bar, 100 μm. **(f)** Whole-body IVIS bioluminescence images on day 21. Mice were injected with equal numbers of CHMp (top) and CHMm (bottom) cells on the mammary fat pads (n = 5 per mouse group). **(g)** Graph showing the total flux of IVIS-imaged mice to measure their growth ability. Quantification of primary tumor size once a week. **(h)** Picture of surgically removed CHMp (top) and CHMm (bottom) primary tumor masses. **(i)** Flow cytometry analysis of CMT cell lines with CD44 and CD24 antibodies. **(j)** Schematic illustration of the isolation of the cancer stem cell population (CD44+/CD24-) in the CIPp cell line. **(k)** An MTT assay was conducted to compare CIPp EVs and CIPp-CSC EVs to analyze CIPm viability. **(l)** The proliferation rate of PBS and EV treatment was determined by manual cell counting. CIPp-CSC EVs decrease the proliferation rate of CIPm cells more than CIPp EVs. All data are presented as the mean ± SEM. Experiments were performed in triplicate if not indicated. The statistical analysis is presented. **P < 0.01 and ***P < 0.001. ns, not significant. Results are presented as mean ± SD (5 mice were used for each group)

It is presumed that the stemness of CSC produced distinct EVs and that the EVs had an antimetastatic growth effect. We further used other primary and metastatic cells for confirmation, CIPp and CIPm cells (Extended Data Fig. 4a), which had fewer differences in CSC features (Fig. 3i), and there was no clear difference between CIPp and CIPm cells in the amount of gene expression associated with ALDH and drug resistance, and stemness transcription factor except Nanog (Extended Data Fig. 4e). Thus, we sorted CIPp cells only into the population of CD44+/CD24-cells, which we named CIPp-CSCs (Fig. 3j). EVs were isolated from the CIPp-CSC portion and added to CIPm cells. CIPp-CSCs inhibited the cell viability (Fig. 3k) and proliferation (Fig. 3l). CIPp-CSC-derived EVs significantly suppressed metastatic cell proliferation compared to CIPp-derived EVs. Collectively, we confirmed that primary tumors with CSC characteristics inhibited the growth of metastases.

### miR-1 is one of the major regulators enriched in pTDEs and suppresses target gene expression in recipient metastases

To determine the miRNAs that play a role in the effects of pTDEs, we performed next-generation sequencing. The procedures for RNA acquisition and QC data from small RNA sequencing are described in Extended Data Fig. 5a-c. In brief, the sequence length distribution of reads showed that the reads were enriched in 17-27 nt and the correlation between each duplicated sample was 0.96 and 0.86, respectively (Extended Data Fig. 5d). The identified novel and known miRNAs are listed in Supplementary Table 2-3. The detected miRNAs corresponding to each chromosome are shown according to the miRDeep2 score (Extended Data Fig. 5e). Chromosome 6 contained the most miRNAs, whereas chromosomes 36 and 38 contained no miRNAs. Detailed bioinformatic analysis equations are described in the Methods section.

We identified 307 and 249 miRNAs from the pTDEs and mTDEs, respectively (Fig. 4a, b). The miRNA involved in cell proliferation was the top GO term in the pTDE miRNA (top). Stem cell regulation was mainly enriched in mTDE miRNAs (bottom) (Extended Data Fig. 5f). Because one miRNA silences multiple genes, we interrogated miRNA target genes using TargetScan and miRDB, which provided a dog reference (Supplementary Table 4). Kyoto Encyclopedia of Genes and Genomes (KEGG) pathway analysis showed that the target genes of pTDE miRNAs were related to cellular proliferation pathways such as cAMP, cGMP-PKG, PI3K-Akt, and the TNF signaling pathway (Extended Data Fig. 5g). Based on the GO and KEGG analyses, we can infer that EV miRNAs may be the cause of the reduced proliferation of metastases by pTDEs. Next, we examined miRNAs that were differentially expressed in pTDEs. Notably, cfa-mir-1-1 and -2 showed the highest expression levels among the miRNAs found in the pTDEs (Fig. 4c). In addition, from the miRNA-gene network analysis, miR-1 showed the highest number of nodes among the miRNAs and the highest degree of miRNA known to play a distinct role in tumor suppression (Fig. 4d).

**Figure 4.**
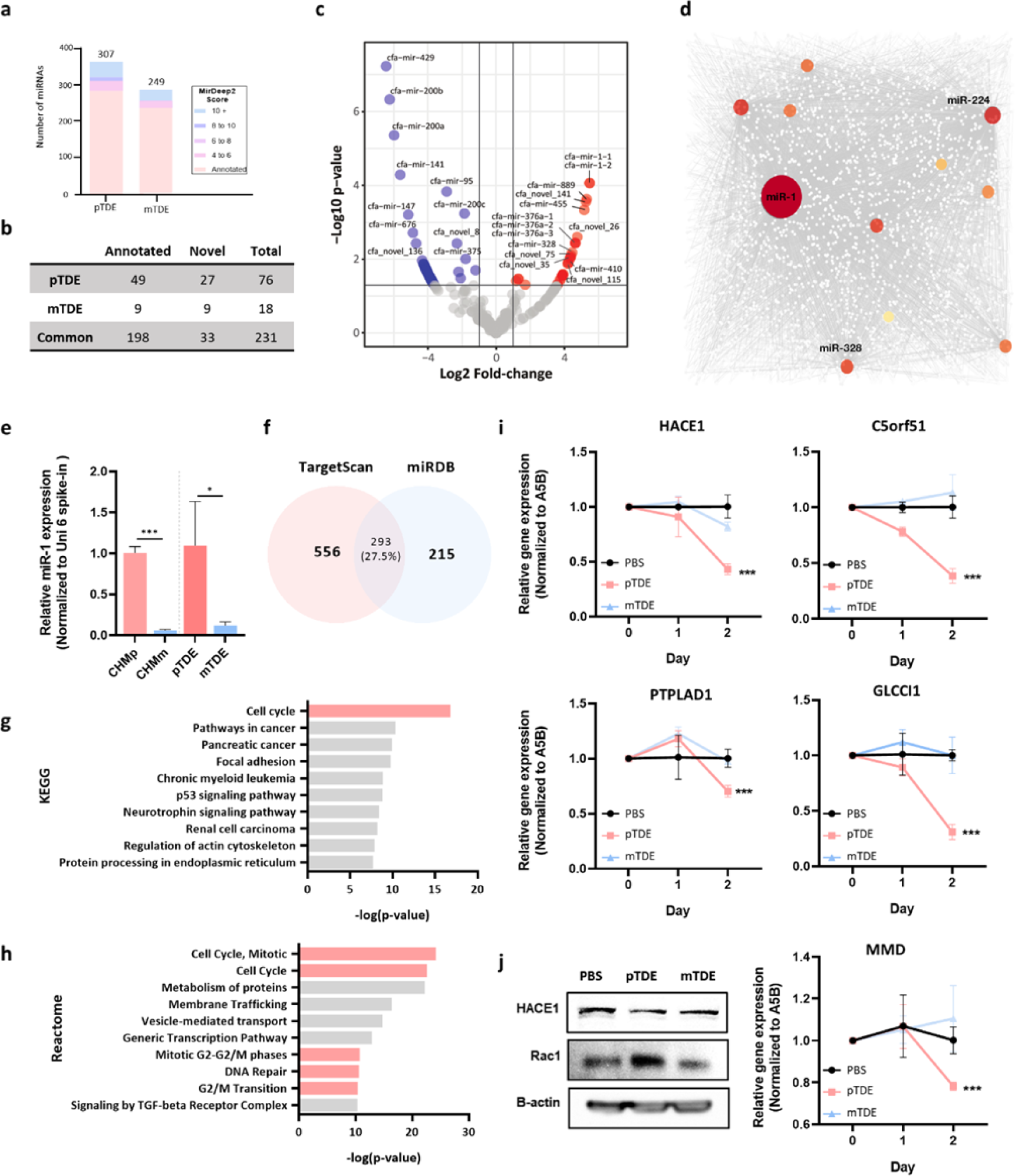
Small RNA sequencing reveals that miR-1 is enriched in pTDEs and that its network has a critical role in cell cycle regulation. **(a-b)** Comparison of the identified miRNAs between pTDEs and mTDEs. **(c)** Volcano plot showing differentially identified miRNAs between pTDEs and mTDEs. Red dots represent pTDE-enriched miRNAs, while blue dots represent mTDE-enriched miRNAs. Gray dots represent miRNAs with -Log10 (p value) and -Log2 (Fold change) values below 1.3 and 1, respectively. **(d)** Network analysis of miRNAs and their target genes was conducted for the top ten pTDE miRNAs. The size of colored dots indicates the number of genes regulated by the miRNA. miR-1 has the largest colored dots, which represent higher interactions than others. **(e)** miR-1 expression between CHMp and CHMm cells at the cellular level (left) and EVs (right). EVs were treated with RNase for 10 min at 37 °C to degrade contamination of EV-free miRNAs. Ct (cycle of threshold) values are normalized to Uni 6 Spike-in within the same cDNA concentration. **(f)** Identification of target genes of miR-1. TargetScan and miRDB were used to screen miR-1 target genes using a dog database. The Venn diagram indicated that TargetScan and miRDB shared 293 genes. **(g-h)** KEGG and Reactome analyses of the top 100 genes shared between TargetScan and miRDB. **(i-j)** Expressional changes in mRNA and protein levels of miR-1 potential target genes. HACE1, C5orf51, PTPLAD1, GLCCI1, and MMD were decreased when pTDEs were administered. Ct (cycle of threshold) values are normalized to A5B gene expression within the same cDNA concentration. The protein levels of HACE1 and the HACE1 target gene Rac1 were analyzed by Western blotting. The statistical analysis is presented. Error bars represent the mean ± SEM. Two-way ANOVA was used to compare groups. *p< 0.05, ***p < 0.001.

Interestingly, the expression pattern of miR-1 between RNase-treated pTDEs and mTDEs corresponded to that in CHMp and CHMm cells (Fig. 4e). We integrated two databases (TargetScan and miRDB) that can retrieve miR-1 target genes and sorted 293 common genes (Fig. 4f). KEGG and Reactome analyses of miR-1 target genes showed significant associations with the cell cycle (Fig. 4g, h). Next, we selected ten miR-1 expected target genes (HACE1, C5orf51, PTPLAD1, GLCCI1, MMD, GJA1, THSB4X, SCAF11, WNK3, and SMIM14) and analyzed whether they were affected by pTDEs via qRT-PCR. Additionally, the levels of HACE1, C5orf51, PTPLAD1, GLCCI1, and MMD were significantly decreased by pTDE treatment in CHMm cells (Fig. 4i). In contrast, the other five genes did not show any significant changes (Extended Data Fig. 6a). Treatment with pTDEs also reduced the protein level of the HACE1 target, which then resulted in an increase in Rac1 protein, a target of the HACE1 E3 ligase ^31, 32^ (Fig. 4j). The regulation of this miR-1-HACE1-Rac1 axis could help explain how pTDEs cause growth inhibition in recipient metastatic cells. Altogether, miR-1 in pTDEs can reduce HACE1 levels in recipient cells and lead to the accumulation of Rac1, which causes elevation of ROS and subsequently induces cell cycle arrest after DNA damage, which slows cell proliferation. Collectively, pTDEs from primary tumors possess miR-1, which might play a role in the suppression of metastasis.

### pTDE-miR-1 has an anti-metastatic effect in an *in vivo* mouse model

To verify that the inhibition of metastatic growth by pTDEs was due to miR-1 present in high amounts, we treated CHMm cells with miR-1 (Extended Data Fig. 7). Treatment with miR-1 encapsulated in liposomes dramatically decreased the expression of six target genes of miR-1 (Extended Data Fig. 7a). Cell viability and growth were also significantly reduced in the group treated with miR-1 (Extended Data Fig. 7b and c). Additionally, similar to conventional pTDEs treatment, miR-1 also enriched the G2/M stage of cells (Extended Data Fig. 7d). Furthermore, miR-1 increased intracellular ROS levels and genomic instability, similar to pTDEs (Extended Data Fig. 7e, f). Lastly, miR-1 increased the levels of γ-H2A and Rac1 while decreasing the levels of HACE1 (Extended Data Fig. 7g). Collectively, miR-1 induced genomic instability, increased G2/M phase, and reduced proliferation, similar to pTDEs.

We then engineered pTDEs with an overload of miR-1 (pTDE-miR-1), resulting in an approximately 500-fold increase in miR-1 concentration compared to that of pTDE (Fig. 5a). pTDE-miR-1 demonstrated significantly enhanced properties compared to pTDEs, including decreased expression of target genes, cell viability, and cell growth, as well as an increase in the G2/M phase, ROS, DNA damage, and dormancy state (Fig. 5b-g, Fig. and Extended Data Fig. 7h, i). These results demonstrate that the growth inhibition of metastases by primary tumors can be exerted by miR-1 present in pTDEs.

**Figure 5.**
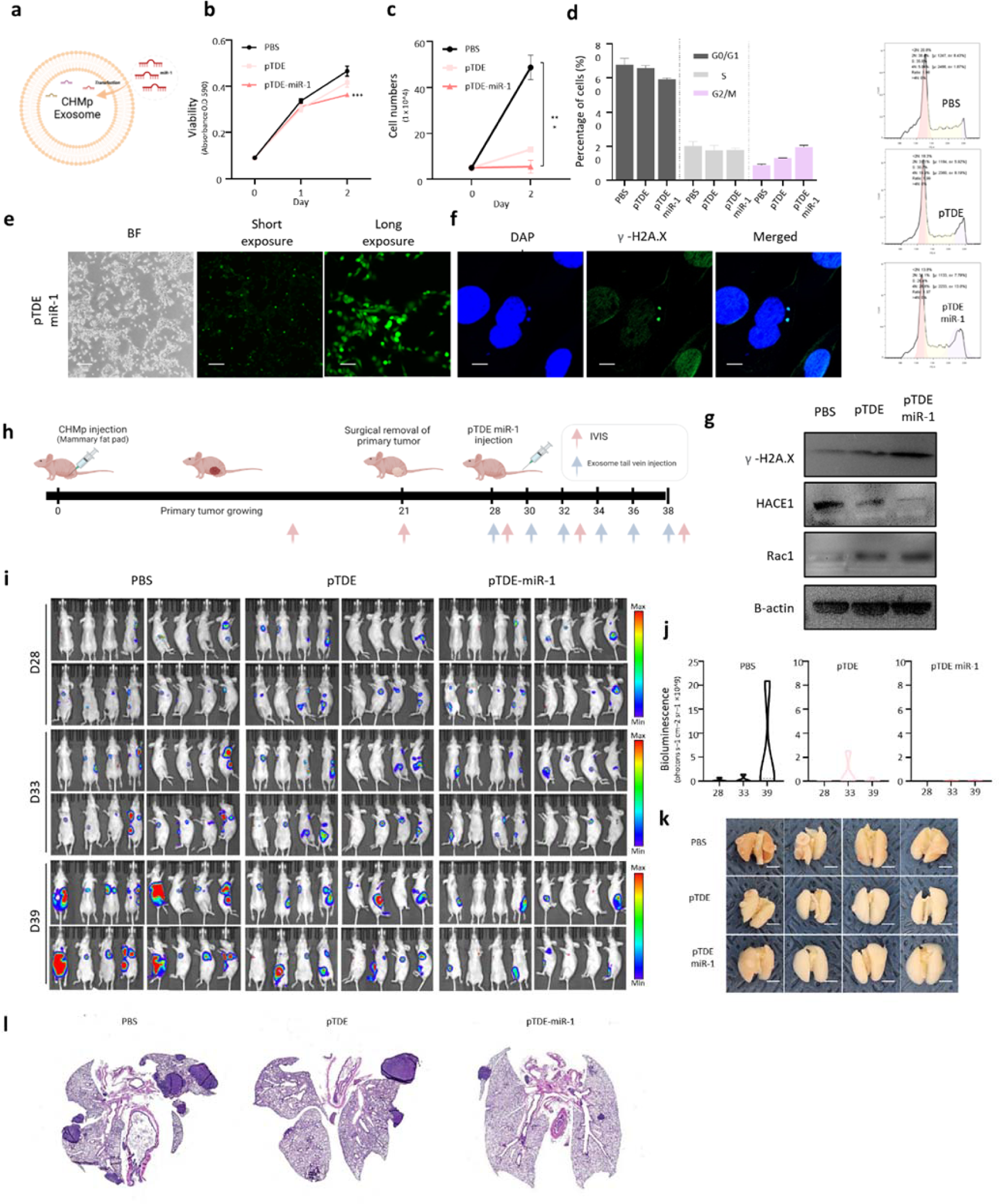
pTDE-miR-1 showed therapeutic effects in mouse model. **(a)** Schematic illustration of miR-1 transfection into pTDEs. **(b-d)** Viability, proliferation and cell cycle assays were conducted in CHMm cells treated with PBS, pTDEs, and pTDE-miR-1. Viability and proliferation were further decreased when cells were treated with pTDE-miR-1 compared with pTDEs. **(D)** pTDE-miR-1 further increases G2/M phases compared with pTDE treatment. (Left) Quantification of cell cycle states in CHMm cells treated with PBS, pTDEs, and pTDE-miR-1. (Right) Representative flow cytometry graph for each treated group. **(e)** Representative images showing cellular ROS levels; brighter green fluorescence indicates high ROS. pTDE-miR-1-treated cells produce higher ROS than pTDE-treated cells. Scale bar, 170 μm. **(f)** For measuring damaged DNA, γ-H2A.X (green) and DAPI (blue) stains were used. The white arrow indicates damaged DNA, which colocalized with γ-H2A.X and DAPI. Scale bar, 10 μm. **(g)** Western blot for H2A.X and miR-1 target genes. Downregulation of HACE1 and upregulation of Rac1 were shown by Western blot. **(h)** Schematic illustration of an established experimental mouse model and injection schedule. Mice were inoculated with CHMp cells into the mammary fat pad to produce a primary tumor. On day 21, primary tumors were surgically removed. Seven days after surgery, mice were treated with PBS (n=4), pTDEs (n=4) and pTDE-miR-1 (n=4). **(i)** Bioluminescence images of lung metastases in mice after treatment with PBS, pTDEs, and pTDE-miR-1 six times at two-day intervals. Bioluminescence images were obtained once a week. **(j)** Graph showing quantification of lung metastases in the mice treated with PBS, pTDEs, and pTDE-miR-1. **(k)** Pictures of mouse lungs after PBS and pTDE treatment. Top; PBS treated, Middle; pTDE treated, and Bottom; pTDE-miR-1 treated. (**l)** Representative H&E staining of lung tissues in the metastasis model. Left two; PBS treated, Middle two; pTDE treated, and Right two; pTDE-miR-1 treated (4X). The statistical analysis is presented. Error bars represent the mean ± SEM. Two-way ANOVA was used to compare groups. *p < 0.05, ***p < 0.001. Results are presented as mean ± SD (4 mice were used for each group)

Next, we examined the effect of pTDE-miR-1 on metastatic growth inhibition in an animal model. As shown in Figure 1, after removing the primary tumor from the CMT mouse model, mice confirmed to have metastases to a secondary organ (lung) were grouped and treated with PBS, pTDEs, or pTDE-miR-1. The injection schedule is shown in Figure 5h. When quantified by radiance, metastatic growth was dramatically suppressed in the pTDE group and mostly in the pTDE-miR-1 group compared to that in the PBS group (Fig. 5i, j). The lungs from PBS-, pTDE-, and pTDE-miR-1-injected mice showed significant differences in metastatic nodule counts (Fig. 5k). Metastases were easily found in mouse lung tissue from the PBS group and scarcely in the pTDE group, and the size of metastatic tumors was smaller in the pTDE group and much smaller in the pTDE-miR-1 group than in the PBS group (Fig. 5l). Overall, these data strongly suggest that EVs derived from primary tumors and loaded with miR-1 can inhibit the formation and growth of metastases that have already formed in lung tissues. These results indicate that pTDEs loaded with miR-1 can be developed to treat metastases by inhibiting their growth.

### Blood EV miR-1 levels can be a diagnostic marker for metastases in clinical specimens

We then investigated whether EV miR-1 could be detected in the blood of patients with CMT or HBC (Fig. 6a). Patient information is described in Supplementary Table 5. EVs were isolated from plasma and serum obtained from the patients and were found to be cup-shaped and less than 150 nm in size (Fig. 6b, c). We observed that the expression of EV-miR-1 was higher in both CMT and HBC patients than in healthy controls; furthermore, its levels increased gradually with the progression of breast cancer (Fig. 6d, e). The largest amount of EV-miR-1 was detected in the sera of Stage III patients, suggesting that primary tumors in locally advanced metastatic stages of breast cancer might generate an increased inhibitory signal of EV-miR-1 to inhibit the growth of veiled metastatic cancer. Next, we investigated the association between miR-1 expression and patient survival using the Kaplan‒Meier plotter. We found that the high miR-1 expression group had better overall survival (OS) in both lymph node-negative and -positive patients, as well as in all specimens. This was expected, as high levels of miR-1 in tumor tissues may lead to high levels of EV-miR-1 in the blood, which could result in better suppression of metastasis. However, the OS patterns of HACE1 differed depending on the metastatic state, as high HACE1 expression was associated with worse OS only in the presence of metastasis (Fig. 6f). These results support the potential use of EV-miR-1 as a diagnostic marker and therapeutic target for both CMT and HBC.

**Figure 6.**
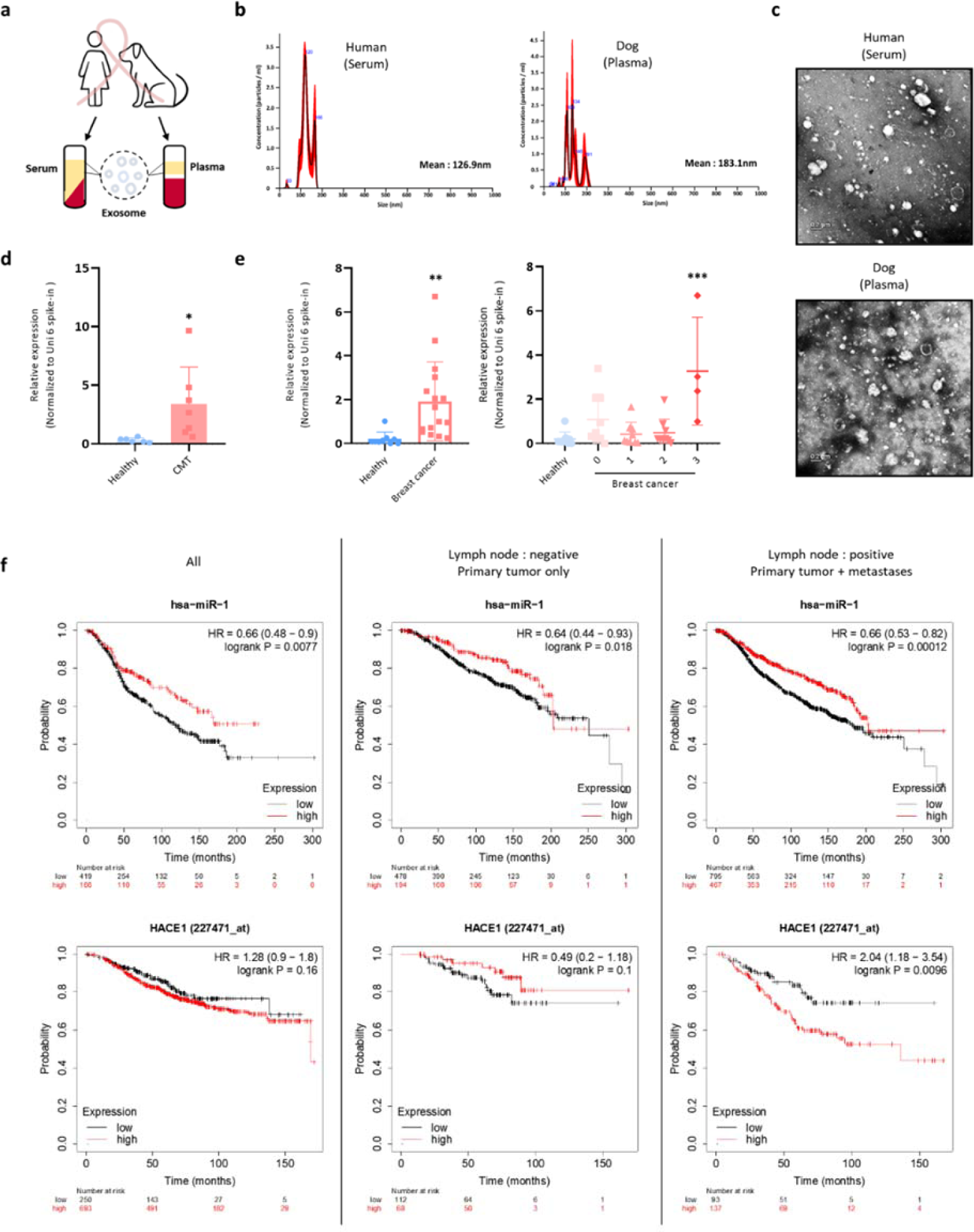
EV miR-1 levels in biological fluids from dog mammary carcinoma and human breast cancer patients. **(a)** Schematic diagram of EVs derived from human breast cancer patients and canine mammary carcinoma patients. **(b)** Nanoparticle tracking analysis (NTA) was used to analyze the size of EVs. Size distribution of human serum-derived EVs (left) and dog plasma-derived EVs (right). **(c)** Transmission electron microscopy (TEM) showed the external appearance of EVs. Top; human serum-derived EVs. Bottom; dog plasma-derived EVs. **(d)** The levels of miR-1 in the EVs of canine mammary carcinoma patients (n=7) and healthy controls (n=6) were examined using qRT‒PCR and normalized to Uni 6 spike-in as a control. **(e)** Expression levels of miR-1 in human breast cancer patients (n=31) and healthy controls (n=9) were examined using qRT‒PCR and normalized with Uni 6 spike-in as the control. miR-1 was higher in breast cancer patients than in healthy controls, especially in late stage (TNM stage III) patients, compared to healthy controls. (stage 0 : n=9, stage Ⅰ : n=9, stage Ⅱ : n =9, stage III : n=4)* p< 0.05, **p < 0.01, ***p < 0.001. **(f)** *In silico* Kaplan‒Meier analysis of breast cancer patients (http://kmplot.com/analysis/). Overall survival (OS) curve comparing the patients with high (red) and low (black) expression of miRNA-1. miR-1 (top) and HACE1 (bottom) expression in breast cancer patients with or without lymph node metastases. Left; analyzed in all patients. Middle: analyzed for lymph node-negative patients. Right: analyzed for lymph node-positive patients.

## Discussion

One of the key questions for unrevealing metastasis is clarifying how the primary tumor controls the cells that exit from their territory. During the initial stages of metastasis, disseminated tumor cells (DTCs) detach from the primary tumor, intravasate into the circulation, and eventually extravasate to colonize distant secondary sites. The emergence of clinically significant metastatic tumors signifies a crucial juncture where previously quiescent DTCs reactivate and regain stem cell-like properties that facilitate self-renewal and the potential for continued tumor growth ^33^. However, the underlying mechanisms governing the transition between dormant and awakened states in these cells remain incompletely understood, including the factors that determine their ultimate fate at secondary sites. Recently, Borriello et al. showed that metastasizing tumor cells are maintained in a dormant state at secondary sites of the lung by primary tumor-associated macrophages^1^. It has been reported that primary tumor can inhibit growth of metastases by inducing apoptosis in DTCs directly or regulating immune cells and endothelial cells in the tumor microenvironment ^1, 3, 15, 34–36^. Because primary tumors inhibit DTCs, when the primary tumor disappears, DTCs start to grow, regardless of their location, due to the release from growth inhibition^37^. This is supported by the fact that early stage surgical removal (on day 4 after initial tumor implantation) of primary breast tumors in mice induces micro-metastases in the lymph nodes because DTCs are present in these nodes at this stage^36^. Conversely, late-stage surgery (on day 13) results in the formation of distant organ metastases, such as the lungs, as DTCs have had sufficient time to disseminate and colonize these organs^38^. Therefore, surgical removal of primary tumors remains a controversial topic in the field of cancer research. While it can improve patient survival and drug accessibility^39^, it can also promote the growth of metastases^40, 41^ ^38^. Here, we revealed that DTCs in a dormant state cannot grow because of EVs derived from primary tumors. We also showed that the lack of EVs due to the absence of primary tumor promotes the growth of DTCs, while primary tumor-derived EVs can inhibit the growth of DTCs, even exhibiting a therapeutic effect.

To demonstrate the mechanism by which primary tumors can inhibit the growth of metastases, we considered that studying the fate of DTCs after removal of primary tumors in mice would be an ideal model. The growth inhibition caused by primary tumor-derived EVs clearly suggests that the primary tumor induces antitumor effects to prevent the development of metastasis (Fig. 1). We demonstrated that this antitumor effect of EVs is caused by the accumulation of ROS and damaged DNA. However, DNA damage resulted in an increase in the G2/M phase of the cell cycle, but not apoptosis. We extended the role of EVs in the tumor mass, similar to endostatin, which has been reported as an anti-angiogenic factor by Folkman ^15^. We confirmed that primary tumor-derived EVs limit the metastatic outgrowth of DTCs by disrupting the endothelial cell formation of capillary-like structures. Consistent with our findings, the clearance of tumor cells in the EV-injected mouse model was mediated by both cellular slow-cycling tumor cells and anti-angiogenesis (Fig. 2). It was also reported that the primary tumor can induce antitumor immunity by priming T cells and macrophages ^1, 3^. Because we used immunodeficient mice, we could not determine the role of EVs in immunity. Further investigation is warranted to determine whether primary tumor-derived EVs behave similarly in immunocompetent mice with intact immune systems.

Nonetheless, we investigated its inhibitory ability on primary tumors and aimed to uncover its characteristics. It is speculated that cells that disseminate from primary tumors and settle in distant locations are less likely to thrive in harsh environments if they lack stemness^43–46^. Therefore, we examined the cancer stemness of the primary tumors to determine their metastatic inhibitory capability. Although many studies have investigated the effect of primary cancer stem cell-derived EVs on DTCs, none have addressed their antitumor effects. Analyses of cancer stemness in primary tumor cell lines CHMp and CIPp revealed that CHMp has cancer stemness (*CD*44^+^/*CD*24^-^/*ALDH*^high^) that approaches 80%, while CIPp, although not to that extent, still has more cancer stemness than its metastatic cell line. Moreover, EVs derived from *CD*44^+^/*CD*24^-^ population of CIPp cells had much higher antitumor activity than EVs derived from the whole CIPp cell population, suggesting that the *CD*44^+^/*CD*24^-^ population is involved in the suppression of metastases (Fig. 3).

Analysis of EV-miRNAs revealed that miRNA-1 is abundantly present in primary tumor-derived EVs, and overloading miR-1 in pTDEs significantly enhances their growth inhibitory effect (Fig. 4-5). Consistent with previous reports that miR-1 inhibits tumor growth and metastasis ^50^, miR-1 is a tumor-suppressive miRNA conserved in humans and dogs that inhibits growth and metastasis in breast cancer ^50^. Furthermore, our study showed that treatment with pTDE resulted in decreased expression of HACE1, a target of miR-1. Decreased expression of HACE1 leads to the accumulation of Rac1, which in turn elevates ROS levels and induces cell cycle arrest. This is consistent with previous studies showing that HACE1 controls ROS generation by catalyzing the ubiquitination of Rac1^31^. Taken together, these findings suggest that miR-1 is involved in the suppression of metastasis. The features of miR-1 expression in primary tumors are not confined solely to CMT, as evidenced by its detection in primary colorectal cancers obtained from human patients ^51^. In human colorectal cancer cell lines, miR-1 expression is also higher in SW480 (primary tumor) than in SW620 cells (metastases) ^52^. In this case, SW480 cells also exhibited a CD133^+^ stem-like phenotype compared to SW620 ^53^. Our findings in dogs provide a valuable basis for research on human patient-derived cell lines, given that miR-1 is conserved in both species. These consistent results between dogs and humans provide evidence of a relationship between primary tumors, cancer stem cells, and miR-1 expression. However, it is noteworthy that EVs contain not only miRNAs but also other nucleic acids, proteins, and lipids, which might play roles in inhibiting metastases.

We observed an intriguing potential for EVs to serve as biomarkers for both HBC and CMT owing to their detectability in biological fluids. Notably, cancer patients with HBC and CMT exhibit increased levels of EV-miR-1, with a progressive elevation observed in HBC patients with advanced TNM stages. We postulate that this may reflect the primary tumor’s augmented release of EV-miR-1 upon DTC dissemination and the colonization of distant organs. Furthermore, in line with our clinical observations, we found that HACE1 gene expression was a predictor of inferior overall survival in patients with both primary tumors and metastases relative to those with primary tumors only (Fig. 6).

In summary, we have provided the first evidence that EVs secreted by cancer stem cells in primary tumors can impede metastatic growth. Moreover, our investigations offer a molecular basis for EV-mediated suppression of DTC proliferation. Specifically, primary tumor-derived EVs harboring miR-1 exert dual effects: inducing ROS-mediated genomic instability and cell cycle arrest in metastatic cells, and impeding angiogenesis in nearby endothelial cells. Furthermore, our findings suggest that EV-miR-1 may hold promise as a diagnostic marker and therapeutic agent for treating metastasis after primary tumor resection.

## Materials and methods

### Cell culture

Six canine mammary gland adenocarcinoma cell lines (CHMp, CHMm, CIPp, CIPm, CTBp, and CTBm) were purchased from N. Sasaki lab (38) and grown in RPMI 1640 medium (HyClone, SH30027) containing 10% fetal bovine serum (FBS; Gibco 1600044) and 50 μg/ml gentamicin (Sigma‒Aldrich, G1272). CHMp and CHMm cells were transfected with a firefly luciferase gene-carrying plasmid (Addgene #18964) using Lipofectamine 3000 reagent (Invitrogen, LM3000015). A culture medium containing geneticin (G418 sulfate) was used to select stably transfected cells (Gibco, 10131035). Twenty-four hours after transfection, G418 (500 µg/ml) was added and maintained for one week to eliminate untransfected cells. A single cell was isolated from stable luciferase-expressing cells to establish a stable cell line and maintained with 250 µg/ml G418 for two weeks. Luciferase expression was confirmed using the luciferase assay. All cells were grown in a humidified incubator at 37 °C with 5% CO_2_ and were confirmed to be negative for mycoplasma contamination.

### Mouse experiments

All mouse experiments were conducted in accordance with the Seoul National University Institutional Animal Care and Use Committee (IACUC) guidelines, and the animal protocol was approved (SNU-210323-1-2). Nude mice (CrTac:NCr-Foxn1nu) were kept in pathogen-free conditions under a 12 h light/dark cycle at a controlled room temperature (22±2 °C). A metastasis model was generated in 5-week-old female nude mice following orthotopic injection of 5 × 10^5^ luciferase-labeled CHMp and CHMm cells into the mammary fat pad, and resection surgery was performed 21 days post-implantation. The mice were anesthetized, and the primary tumors were resected. Mice that underwent surgery were monitored for symptoms of pain and sacrificed by carbon dioxide (CO_2_) inhalation. IVIS bioluminescence imaging was performed during the development of spontaneous lung metastases. All mice were randomized before the injection of EVs and blindly selected before injection. For EV injection, all treatments were administered intravenously in a final volume of 150 µL. The mice were treated with EVs six times at two-day intervals. The experimental endpoint followed the IACUC guidelines, and the maximal tumor volume was never exceeded.

### EV isolation and labeling

EVs were purified from cells cultured under serum-free conditions by using a combination of ultrafiltration and ultracentrifugation. The graphical method is illustrated in Supplementary figure 2a. Cells were grown to 80% confluency, washed twice with PBS, and incubated in serum-free medium for 24 h. First, the cell culture supernatant was subjected to a differential centrifuge to eliminate cells, dead cells, and cell debris, and filtered sequentially with 0.45 and 0.22 μm filters. The filtered supernatant was concentrated using 10 K Amicon-Ultra 15 Centrifugal Filter Units (Merck, UFC903024). The filtered units were then centrifuged sequentially. ultracentrifugation for 80 min and washing with PBS. The EV concentration (0.1ug/ml was applied in all *in vitro* assays.

EVs from biological fluids were isolated using ExoQuick exosome precipitation solution (System Biosciences, SBI-EXOQ5A-1). Plasma and serum samples were centrifuged at 3,000 × g for 15 min to remove cells and cell debris. ExoQuick was then added to the supernatant at an appropriate volume and incubated for 30 min at 4 °C. Pelleted EVs were resuspended in Qiazol (Qiagen, 79306) for RNA isolation and urea/SDS lysis buffer for protein isolation. Isolated EVs were stored at −80 °C for later use.

EVs were labeled with PKH67 lipophilic membrane dye (Sigma, MNI67-KIT), following the manufacturer’s instructions. Briefly, isolated EVs were resuspended in 1 ml of Diluent C and 6 µL of PKH67 dye was added. The mixture was incubated for 5 min at room temperature and centrifuged at 100,000 g for 80 min.

### Nanoparticle tracking analysis (NTA)

NTA was used to characterize the size and concentration of EVs in the cell culture supernatant and biological fluids using the NanoSight LM10 model (Malvern). The samples were diluted with PBS (0.22 µm filtered) and injected into the laser chamber. The data were analyzed using NTA v3.2 software.

### Transmission electron microscopy (TEM)

The morphology of EVs was analyzed by TEM using Talos L120C (Czech Republic). Briefly, the samples were stained using a negative staining method with 2% uranyl acetate. One drop of diluted EVs was placed onto a glow-discharged copper/carbon-coated grid. After 1 min, the grid was drained using filter paper, and a drop of 2% uranyl acetate was added. The staining solution was drained, and the samples were observed using TEM (120 kV).

### Small RNA sequencing and data analysis

The miRNA-seq library was prepared using the Small RNA Library Prep Kit (Nextflex) and sequenced as 100 bp or 150 bp paired-end reads on the Illumina HiSeq 3,000 and NovaSeq 6,000 platforms. To remove adapters with low-quality reads and extract miRNA-specific sequences, cutadapt was used with options (--quality-base 33–u 4–m 22–M 30–f fastq–q 20–O 6–j 23–adapter sequence). In this step, two different adapter sequences (TGGAATTCTCGGGTGCCAAGG and GATCGTCGGACTGTAGAAC-TCTGAAC) were used for the forward and reverse reads in paired-end sequencing. Because the trimmed reads were short (22–30 bp), forward and reverse reads in the same sample were merged into one fastq-formatted file. Before and after trimming, the quality of the sequenced reads was estimated using FastQC.

For known and novel miRNA analysis, the miRDeep2 package was used. Before analysis, the following two files were prepared:1) sequence files of mature and hairpin forms of dog miRNAs, which were downloaded from the miRbase database and extracted using the extract_miRNAs.pl script, and 2) indexed files from the dog reference genome (CanFam3.1) using bowtie. First, all filtered read data were merged into a single file for the novel miRNA analysis. The merged data were converted to a collapsed fasta-formatted file and aligned to the reference genome using the mapper.pl script with the options (-e, h, j, m, p). Second, novel miRNAs were identified using the miRDeep2.pl script. The identified mature and hairpin forms of novel miRNAs were extracted and combined with previously prepared known forms of miRBase. Finally, the expression values of known and novel miRNAs were calculated using Quantifier. pl script. The counts per million (CPM), which is counts scaled by the total number of reads, was used for further analysis. MiRDeep2 analysis provides an miRNA score ranging from −10 to 10, with a higher score representing a genuine miRNA. We set a cutoff of 4 for the strict identification of novel miRNAs. To estimate data reproducibility between replicate samples, Pearson correlation values were calculated and visualized using the correlation function in R. For differentially expressed miRNA analysis, fold-change in expression and significance (p-value) were calculated using the EdgeR package in R. Using these calculated values, a volcano plot was visualized through the ggplot package in R.

### miR-1 mimic transfection

Synthetic miRNA mimics have been used to overexpress and overload microRNAs into cells and EVs. CHMm cells were transfected using the lipid carrier Lipofectamine RNAiMAX (Invitrogen, 13778150), following the manufacturer’s instructions. Fifty picomoles of miR-1 mimics were mixed with the RNAiMAX reagent and incubated for 15 min at RT. miR-1 and RNAiMAX complexes were added to the cells and incubated for 24 h at 37 °C in a CO_2_ incubator. EVs were transfected using 0.3 M CaCl_2_ following the modified CaCl_2_-mediated transfection method ^58^. Forty micrograms of EVs were mixed with 100 pmol of miR-1 mimics in BPS with 0.3 M CaCl_2_ and incubated on ice for 30 min. The mixture was then heat-shocked at 42 °C for 60 s and incubated on ice for 5 min. Transfected EVs were isolated by ultracentrifugation and washed with PBS.

### Clinical specimens

All study protocols and specimen collection were approved by the Institutional Review Board (IRB) of Seoul National University (IRB#SNU 16-10-063) and were performed in accordance with the guidelines. Informed consent for specimen collection was obtained from all subjects, including both human and dog guardians, when they were enrolled.

### Statistical analysis

Data are presented as the mean ± *SEM*. Statistical analysis was performed using the Prism software (v.8.0.1, GraphPad Software). For the statistical significance of multiple groups, 2-way ANOVA combined with Tukey’s honest significant difference (HSD) test was performed, and Student’s t-test was performed for analysis between CHMp and CHMm cells. Significant differences are indicated by different letters (*p<0.05, **p<0.01, and ***p<0.001) in each figure legend. The number of experimental repeats and the value of n are indicated in the figure legend.

## Acknowledgments

We thank Dr. Wan-Hee Kim, College of Veterinary Medicine, Seoul National University, and Dr. Sun-Young Hwang, Haemaru Referral Animal Hospital, for providing valuable canine clinical specimens. We also thank Dr. Jeong Eon Lee and Soo-Youn Lee at Samsung Medical Center for their assistance in providing valuable human clinical specimens.

## Author contributions

Writing – original draft: HSK

Writing – review & editing: JYC and KHL

Methodology: HSK, KHL

Mouse experimental methodology: HSK and TJS

Bioinformatics methodology: KHS

Visualization: HSK and KHS

Conceptualization and study supervision: JYC

## Data and materials availability

All raw and processed EV miRNA-seq data for the CHMp and CHMm cell lines and their biological replicates generated in this study have been deposited in the NCBI Gene Expression Omnibus (GEO) database under accession number GSE213969.

## Code availability

The main code scripts used for miRNA-seq processing and visualization are described in detail and are available at GitHub: https://github.com/snu-cdrc/exosomal-miRNA

## Extended data

**Extended Data Figure 1.**
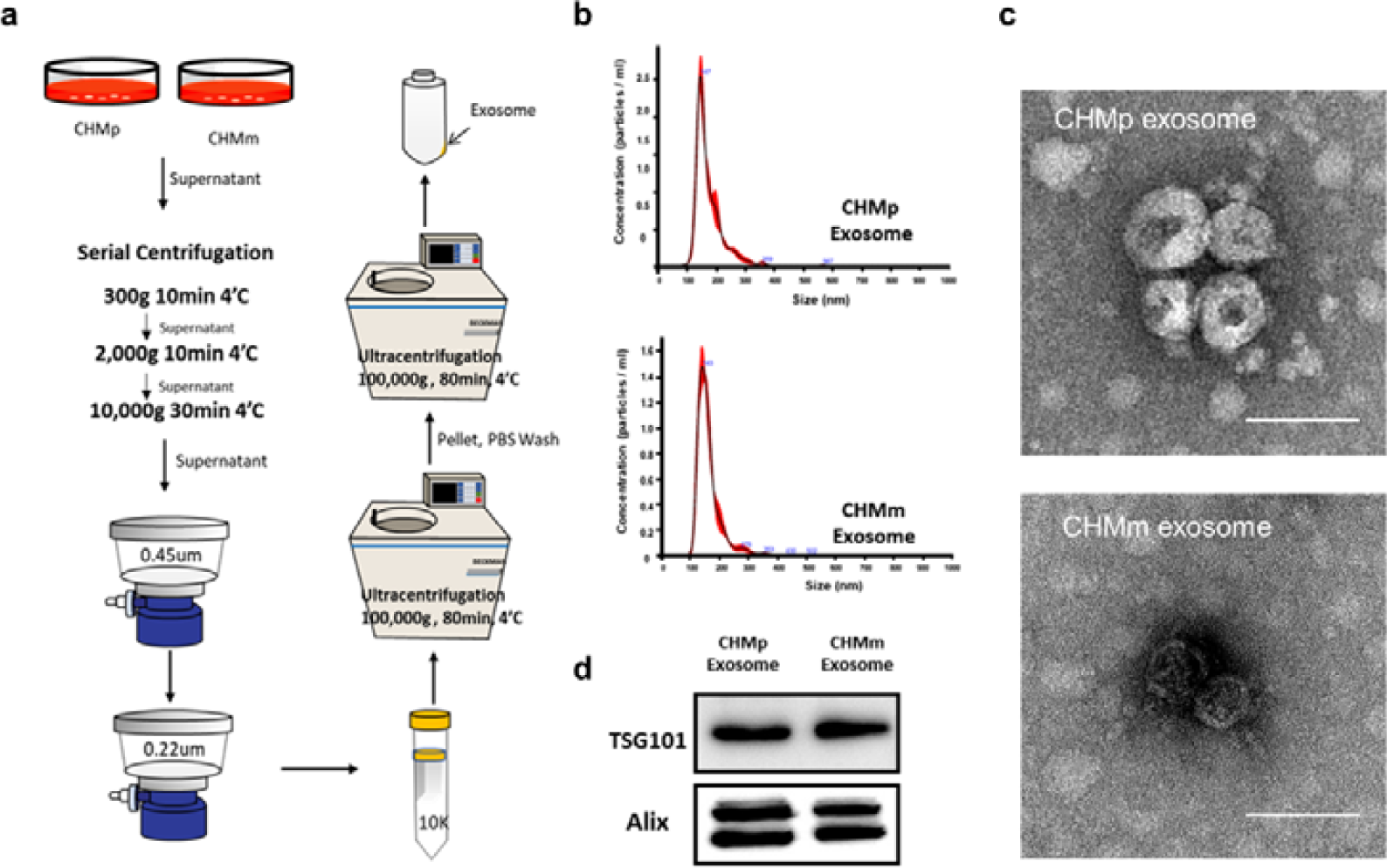
Isolation and characterization of EVs. **(a)** Schematic illustration of EV isolation. Differential centrifugation, ultracentrifugation, and ultracentrifugation were sequentially performed. **(b)** Nanoparticle tracking analysis (NTA) was used to analyze the size of EVs. Size distribution of CHMp EVs (pTDEs, top) and CHMm EVs (mTDEs, bottom). **(c)** Transmission electron microscopy (TEM) showing the external appearance of the EVs. Top: CHMp EVs (pTDEs); Bottom: CHMm EVs (mDTEs). Scale bar = 100 nm, and white arrows indicate cup-shaped EVs. **(d)** EV markers TST101 and Alix were identified using western blotting.

**Extended Data Figure 2.**
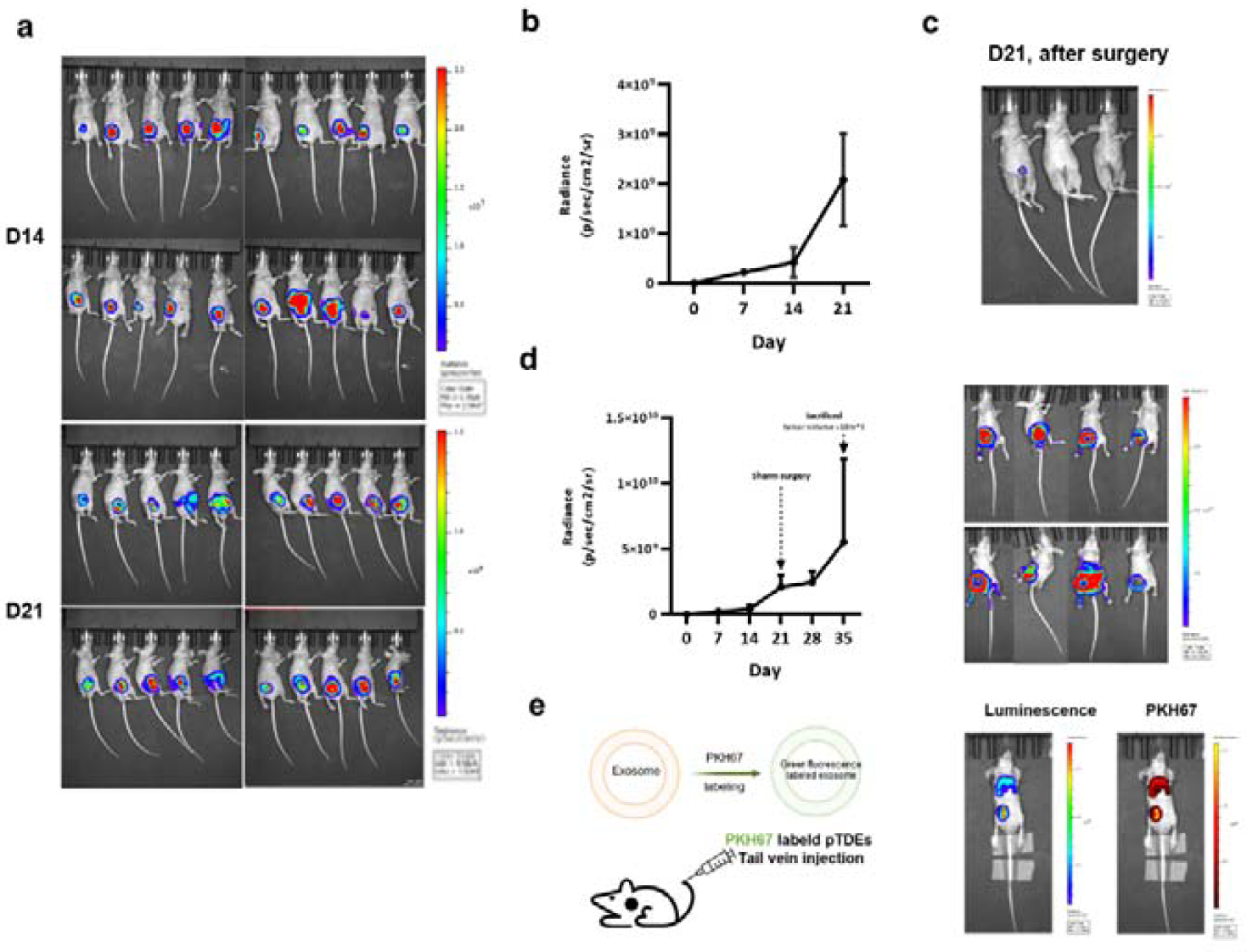
Establishment of a metastasis mouse model. **(a)** Sequential bioluminescence images of primary tumor-growing nude mice from days 14 to 21 using IVIS. **(b)** The bioluminescence intensity was analyzed using an IVIS imaging system. Over time, an increase in the total flux intensity indicated the growth of the primary tumor (imaged once a week, n=20). **(c)** Immediately after surgery, we confirmed that the primary tumors were not visible by IVIS, nor were metastases in secondary organs, such as the frequently metastasized lungs. Representative bioluminescence images of surgically resected primary tumors in mice. D21 bioluminescence images were acquired immediately after the surgery. Left: primary tumor resident mouse, Middle & Right: primary tumor completely resected mouse. **(d)** Quantification of bioluminescence intensity and representative images of the sham group representing tumor growth (n=4). The dashed arrows indicate the day of surgery. Top: Day 28, Bottom: Day 35 **(e)** Establishment of the PKH67-labeled EV injection model. PKH67-labeled EVs were injected into mice via the tail vein and monitored using IVIS imaging. PKH67-labeled EVs accumulated at the primary site and at metastases 48 h after injection.

**Extended Data Figure 3.**
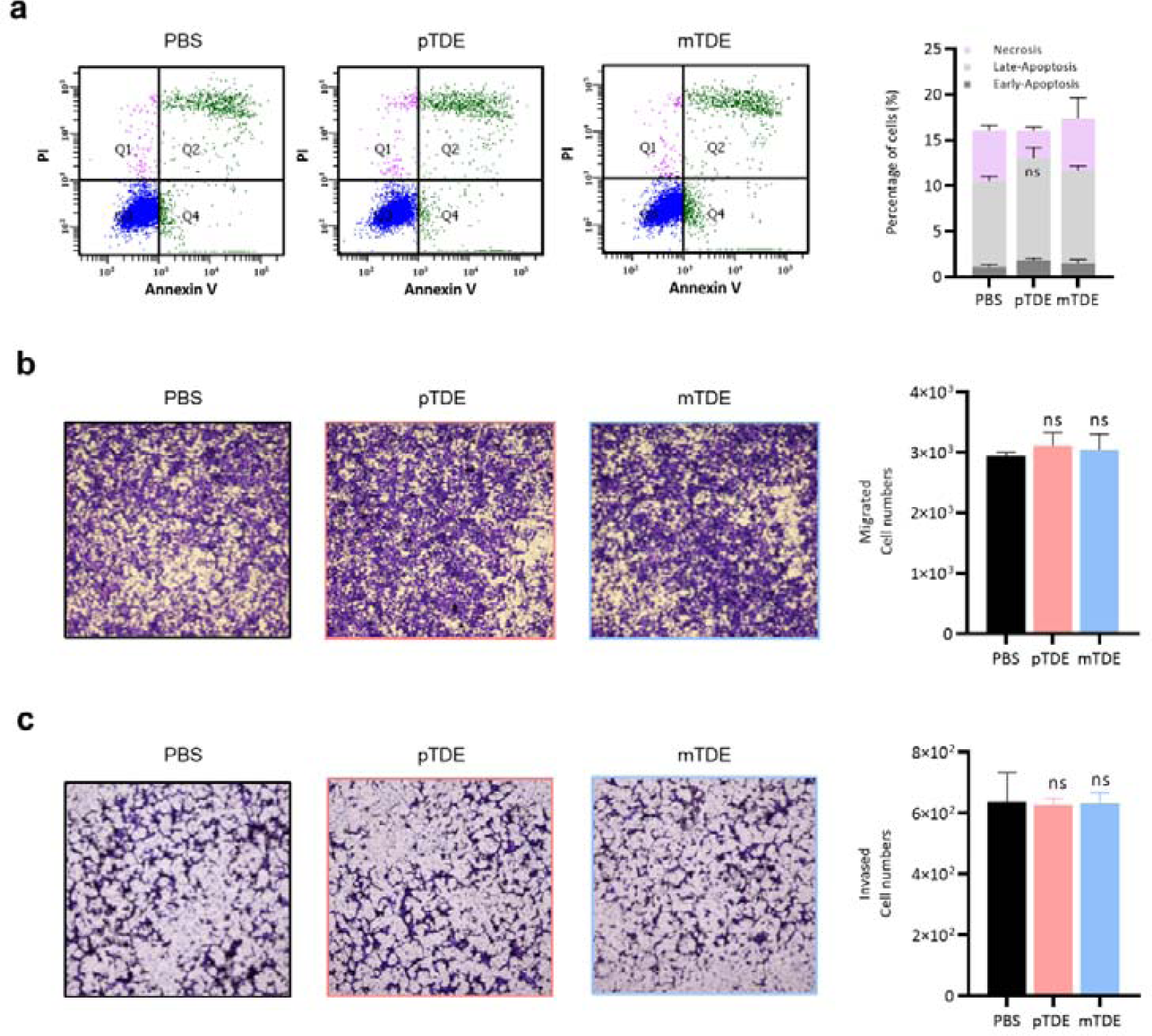
pTDEs are not related to apoptosis or cell mobility. **(a)** EV-treated cells were stained with annexin V and PI to detect apoptosis. The upper left (necrosis), upper right (late apoptosis), and lower right (early apoptosis) sections were quantified by flow cytometry. **(b)** Transwell migration assay using EV-treated cells. Representative images of migrated cells were fixed and stained with crystal violet. The rate of migration was analyzed using the ImageJ software for quantification. No clear differences were observed between the groups. **(c)** Transwell invasion assay of EV-treated cells. Representative images of matrigel-invaded cells were fixed and stained with crystal violet. The rate of invasion was analyzed using the ImageJ software for quantification. No clear differences were observed between the groups. All data are presented as mean ± SEM (n=3). Two-way ANOVA and Tukey’s HSD test were used to compare groups; **P < 0.01, *** p < 0.001, and ns, not significant. Results are presented as the mean ± SD.

**Extended Data Figure 4.**
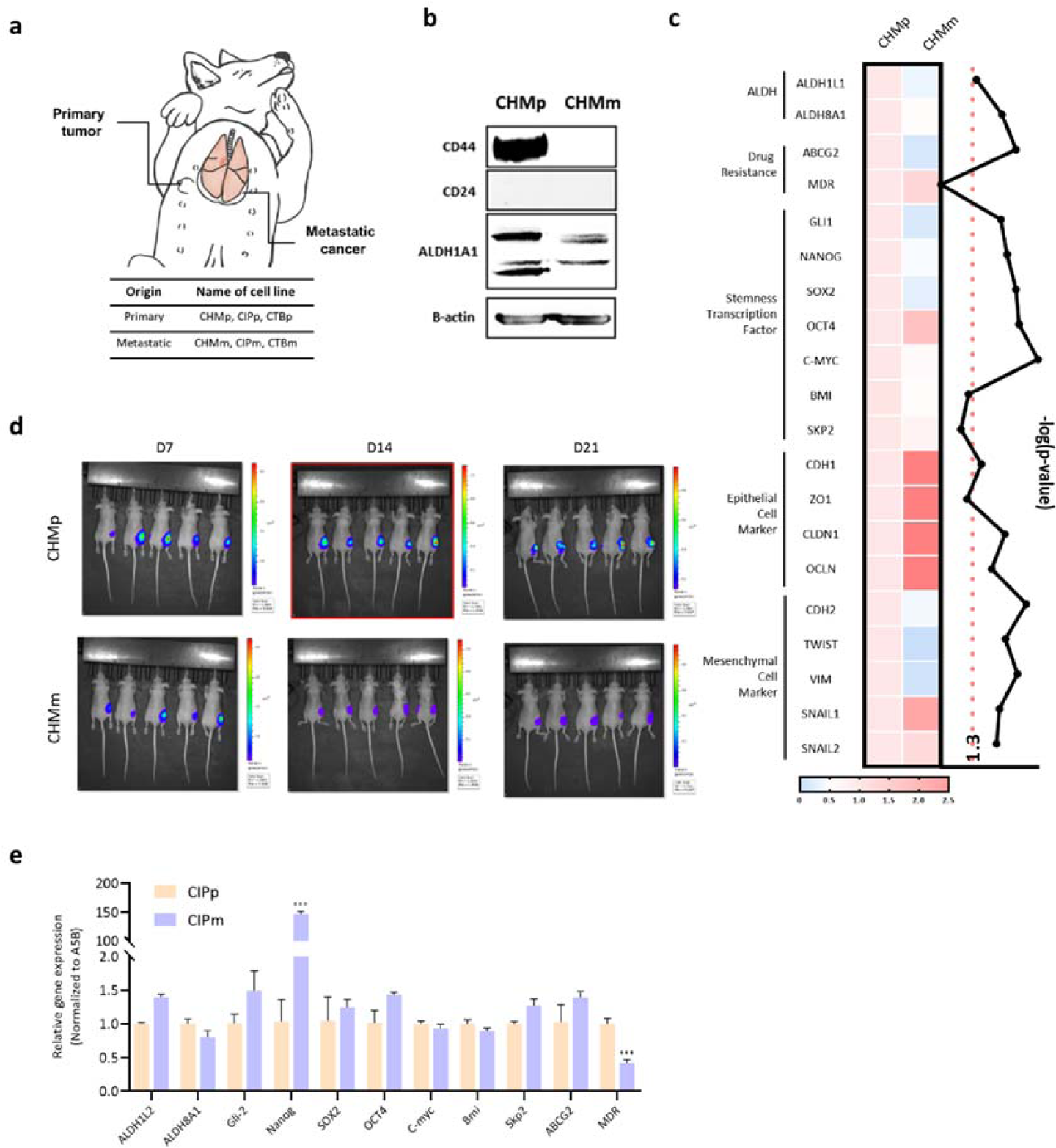
Primary tumor cell possesses more cancer stemness than metastases. **(a)** Graphical illustration of the primary tumor and metastases of canine mammary carcinoma. **(b)** Western blotting for CD44, CD24, ALDH1A1, and β-Actin. CD44 and ALDH1A1 expression significantly increased in CHMp cells. β-Actin was used as an internal control. **(c)** Relative mRNA expression of ALDH, drug resistance, stemness transcription factor, epithelial cell, and mesenchymal cell markers was quantified by qRT-PCR for CHMp and CHMm cells. The results were analyzed using the 2−ΔΔCT method, with A5B as a reference gene. The color bar indicates fold change compared to CHMp cells. Statistical significance between cells was assessed using Student’s t-test. The back line indicates the log(p-value). The red dotted line indicates p < 0.005. **(d)** Sequential bioluminescence images of nude mice injected with CHMp and CHMm cells from days 7 to 21 using IVIS. **(e)** Gene expression of cancer stemness characteristics in CIPp and CIPm cells. ALDH (ALDH1L2 and ALDH8A1), stemness factor (Gli-2, Nanog, SOX2, OCT4, C-myc, and Bmi), and drug resistance (ABCG2 and MDR) genes were used for qRT-PCR. All data are presented as mean ± SEM (n=3). The Student’s t-test was used to analyze the data, and statistical analysis was performed. **P < 0.01 and ***P < 0.001.

**Extended Data Figure 5.**
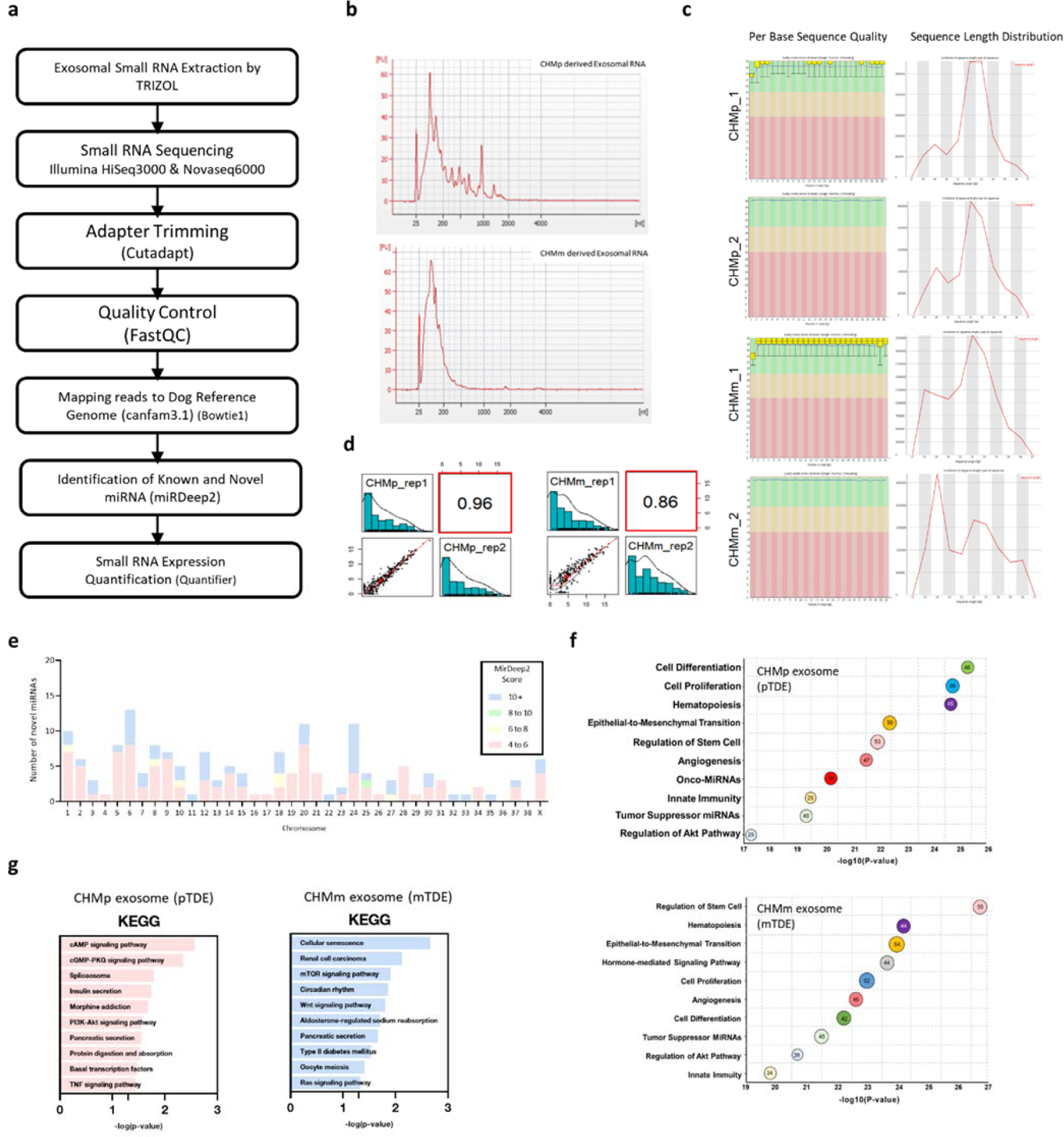
Small RNA sequencing of primary and metastasis-derived EVs. **(a)** Flow chart of EV miRNA sequencing procedure. **(b)** Bioanalyzer electropherograms of the isolated CHMp (top) and CHMm (bottom) EV miRNAs. **(c)** Per base sequence quality (left) and sequence length distribution (right) of the EV miRNA sequencing dataset. Top two; CHMp replicates (n=2), Bottom two; CHMm replicates (n=2). **(d)** Pearson correlation between EV miRNA sequencing replicates. Left: CHMp replicates (n=2), Right: CHMm replicates (n=2). **(e)** Novel miRNAs identified by the miRDeep2 score were distributed across canine chromosomes. **(f-g)** Small RNA expression profiles of primary and metastasis-derived EVs. **(f)** Comparison of miRNA expression between CHMp EVs (pTDEs) and CHMm EVs (mTDEs). **(g)** Differential expression of miRNAs between pTDEs and mTDEs was functionally enriched using TAM2.0. The size of the bubble indicates the number of genes involved. KEGG analysis of pTDE miRNAs (top) and mTDE miRNAs (bottom).

**Extended Data Figure 6.**
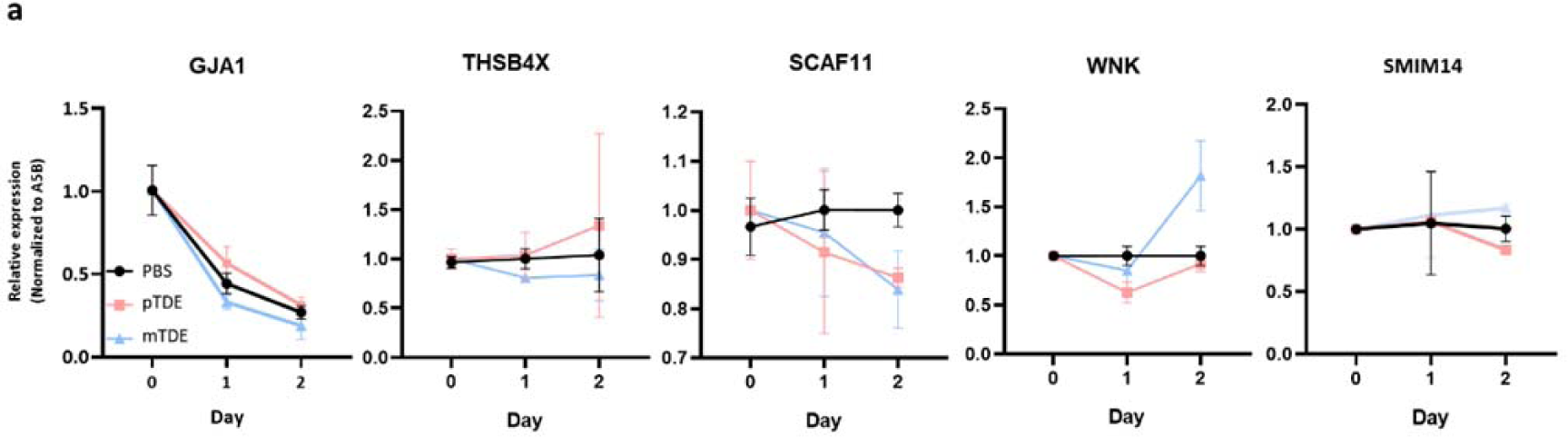
Expression profiles of miR-1 target genes. **(a)** Endogenous mRNA expression of miR-1 target genes in CHMm cells after EV treatment. All data are presented as mean ± SEM (n=3). Unpaired Student’s t-test and two-way ANOVA were used to compare the groups. **P < 0.01 and ***P < 0.001. ns not significant.

**Extended Data Figure 7.**
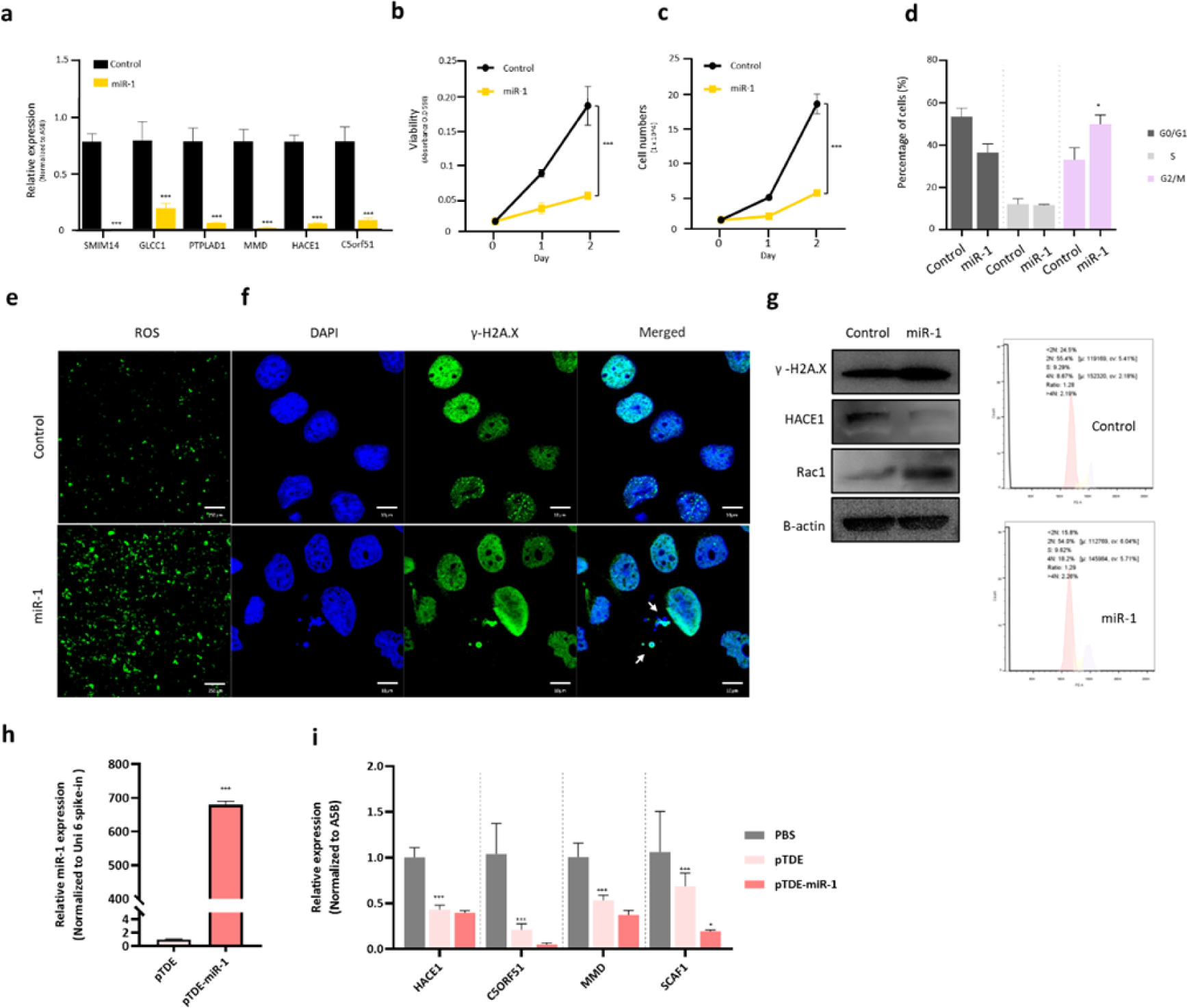
miR-1 transfection into CHMm cells and pTDEs. **(a-d)** miR-1 overexpression in CHMm cells showed similar effects to pTDE treatment. miR-1-transfected cells express low levels of miR-1 target genes (a). miR-1 decreased viability (b) and proliferation rate (c), and increased G2/M phase cell cycle arrest (d) in CHMm cells. (d) (Top) Quantification of the cell cycle states in CHMm cells treated with miR-1. (Bottom) Representative flow cytometry graphs of each treatment group. **(e)** Representative images showing the cellular ROS levels. Brighter green fluorescence indicates high ROS levels. Scale bar, 250 µm. **(f)** To measure DNA damage, γ-H2A.X (green) and DAPI (blue) stains were used. The white arrow indicates damaged DNA that colocalizes with γ-H2A.X and DAPI. Scale bar, 10 µm. **(g)** western blotting for γ-H2A.X and miR-1 target genes. Downregulation of HACE1 and upregulation of Rac1 were observed using western blotting. **(h)** Increased miR-1 levels in pTDE-miR-1 compared with pTDEs. Uni 6 spike-in was used as the control. **(i)** Changes in the mRNA levels of potential miR-1 target genes. C5orf51, MMD, and SCAF1 levels decreased with pTDE-miR-1 treatment. Ct (cycle of threshold) values are normalized to A5B gene expression within the same cDNA concentration. The unpaired Student’s t-test was used to compare the groups. (\**p <* 0.05, \*\*\**p <* 0.001).

## Supplementary Tables

**Supplementary Table 1. Information on the PCR primers and antibodies used**

**Supplementary Table 2. Identified EV miRNA lists**

**Supplementary Table 3. Identified novel miRNA lists**

**Supplementary Table 4. miRNA-mRNA interaction gene lists**

**Supplementary Table 5. Dog MGT and human BC patient information**

